# A novel, wave-shaped profile of germline selection of pathogenic mtDNA mutations is discovered by bypassing a classical statistical bias

**DOI:** 10.1101/2023.11.21.568140

**Authors:** Melissa Franco, Auden Cote-L’Heureux, Zoë Fleischmann, Zhibin Chen, Mark Khrapko, Benjamin Vyshedskiy, Maxim Braverman, Konstantin Popadin, Sarah Pickett, Dori C. Woods, Jonathan L. Tilly, Douglas Turnbull, Konstantin Khrapko

**Affiliations:** Department of Biology Northeastern University, Boston, Massachusetts, USA; Cambridge University, Cambridge, UK; Department of Mathematics, Northeastern University, Boston, Massachusetts, USA; School of Life Sciences, École Polytechnique Fédérale de Lausanne; Swiss Institute of Bioinformatics, Lausanne, Switzerland; Center for Mitochondrial Functional Genomics, IImmanuel Kant Baltic Federal University, Kaliningrad, Russia; Wellcome Centre for Mitochondrial Research, Translational and Clinical Research Institute, Newcastle University, Newcastle upon Tyne, UK

## Abstract

The shift of the level of disease-causing mtDNA mutations (heteroplasmy) from mother to child is typically negatively correlated with the mother’s heteroplasmy (H_m_). In other words, mothers with low H_m_ tend to have children with a higher mutation level (H_ch_) than their own. In contrast, mothers with high H_m_ typically see a decrease in heteroplasmy in their children. This trend has been commonly interpreted as a result of a descending germline selection profile, i.e., positive selection at low H_m_, gradually turning negative at high H_m_. Here we demonstrate, however, that the negative correlation is mostly driven by RTM, or ‘Regression To the Mean’, a classical statistical bias. We further show that RTM can be nullified by using the average between the mother’s and child’s heteroplasmy, as a new variable, instead of the commonly used mother’s heteroplasmy in blood. Additionally, we demonstrate that mother/child average is a better approximation of the actual *germline* heteroplasmy. Moreover, the elimination of RTM revealed a previously hidden wave-shaped HS-profile (positive mother-to-child shift at intermediate average mother-child heteroplasmy, decreasing towards high and low average heteroplasmy). In confirmation of this finding, we show that simulations that involve both wave-shaped HS-profile and RTM, reproduce the observed patterns of inheritance of mtDNA mutations in unprecedented detail. From the health care perspective, the uncovering of the wave-shaped HS-profile (and the removal of the RTM bias) are crucial for families affected by mtDNA disease. From the fundamental perspective, the wave- shaped profile offers a novel understanding of the dynamics of mtDNA in the germline and a novel potential mechanism that prevents the spread of detrimental mtDNA mutations in the population.

**Significance:** From the clinical perspective, the existence of wave-shaped selection may improve predictions and decisions for families affected by mtDNA diseases. From the fundamental perspective, it provides insight into the dynamics of general mtDNA mutations in the germline and in the population, as long as they follow wave-shaped selection profile. In **Fig. 1**, blue and red arrows represent the direction of expected changes of the heteroplasmy in a lineage with time/generations. With wave-shaped selection (**Fig. 1B**), a great majority of nascent low-fraction mutations are expected to converge back to zero and vanish. However, due to random intracellular genetic drift, some mutations will, occasionally, expand and enter the range of positive selection. Then they will be expanded by the selection to higher, detrimental levels, and become prone to downstream removal via death of highly mutated germ cells or inability of highly sick individuals to continue their lineage. In this way, the wave-shaped selection may help to prevent the spread of detrimental mutations in the population and in the species. In contrast, if the descending selection profile (**Fig. 1A**) was in effect, the nascent low heteroplasmy detrimental mutations would have been pushed to intermediate heteroplasmy levels where they will stay longer in ‘hidden disease carriers’ enabling effective spread of mutation in the population.

---

**Fig. 1A**, illustrates this trend is for the most prevalent pathogenic mtDNA mutation, 3243A>G, causing MELAS (Mitochondrial encephalomyopathy, lactic acidosis and stroke- like episodes). Each dot represents a heteroplasmy shift in a mother/child pair, plotted as a function of mother’s heteroplasmy (H_m_). The bold descending “HS profile” (grey regression line) is commonly (and intuitively) explained by the systematic enrichment of the mutation in or among the germline cells (“germline selection”). Thus we typically assume positive selection at low H_m_, gradually decreasing and turning negative at high H_m_ (**Suppl. Note 1**). Here we demonstrate, however, that this intuitive interpretation is totally incorrect: in fact, the decrease of HS is mostly driven by the classical statistical bias called Regression To the Mean (RTM). RTM, in turn, results from random variance in the data generated presumably by random processes in the somatic and germline cells mtDNA and measurement error. We further show that RTM can be nullified by using the average between the mother and child’s heteroplasmy, (H_m_+H_ch_)/2 as the variable, instead of the conventionally used H_m_. Additionally, we argue that (H_m_+H_ch)_/2 is a better approximation of the *germline* heteroplasmy than either H_m_ or H_ch_ (which are measured in a somatic tissue (blood)).

**Figure 1.**
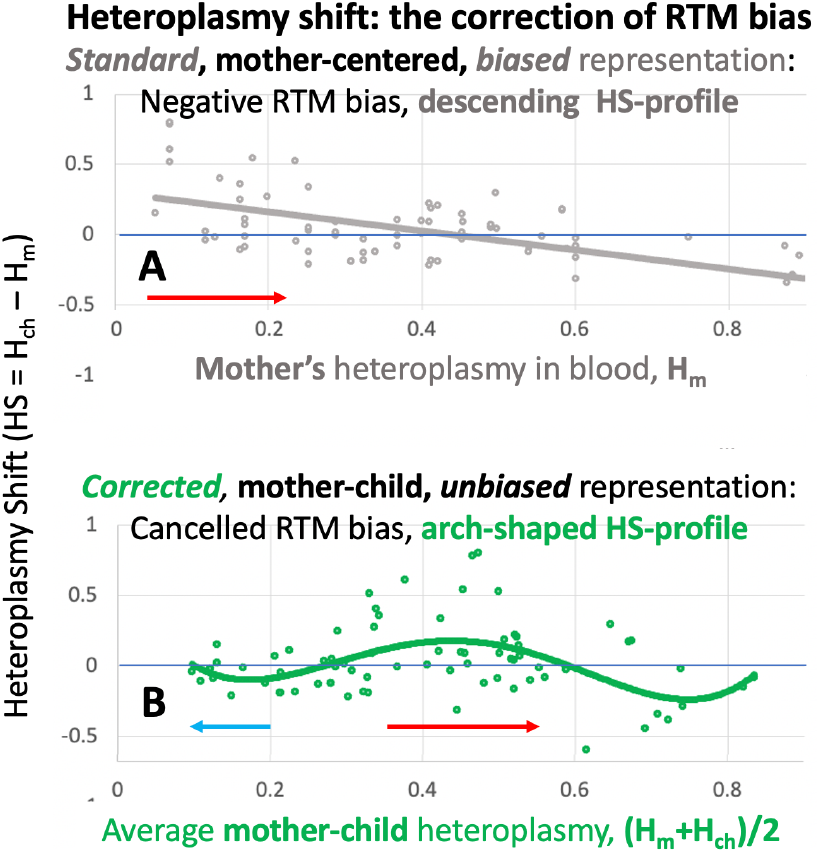
(Graphic Summary) Pathogenic mtDNA mutations that cause a host of devastating inherited diseases are usually thought to follow an intriguing inheritance trend: Mothers with low levels of mutation (called mother heteroplasmy, H_m_) tend to bear children with higher child heteroplasmy (H_ch_) then their own which constitutes positive Heteroplasmy Shift (HS=H_ch_-H_m_). In contrast, mothers with high heteroplasmy H_m_ bear children with lower heteroplasmy H_c_ (negative HS).

**Fig. 1B** shows that the use of (H_m_+H_ch_)/2 removes RTM bias – there is no overall negative trend. Intriguingly, this also revealed a novel, wave-shaped HS-profile (green regression curve). We then used simulations to confirm that a combination of RTM bias and wave-shaped selection indeed led to the observed mother-child inheritance data patterns, including some fine detail. We further show that a few other pathogenic mtDNA mutations, when corrected for the RTM bias by the use of the (H_m_+H_ch_)/2 variable, also exhibit wave-shaped germline selection patterns.

## Introduction

mtDNA is present in many copies per cell, so mtDNA mutations are uniquely positioned for effective selection within cells, including germ cells. Also, because mitochondria are intimately involved in cell death and proliferation, mtDNA mutations are poised to affect germ cell attrition and proliferation. Collectively, these processes are thought to result in germline selection which has a major impact on the transmission of devastating mtDNA-based diseases, and, more generally, on the penetration of detrimental mtDNA mutations into populations and species. Germline selection has been actively discussed as a key means of eradicating of detrimental mutations since it was demonstrated that mtDNA mutations may be subject to negative germline selection (Fan et al. 2008; Stewart et al. 2008). On the other hand, the persistence of certain disease-causing mtDNA mutations in the human population hinted that detrimental mtDNA mutations may be, counterintuitively, under *positive* selection, as often proxied by the heteroplasmy shift (HS), i.e. the difference between the child’s and the mother’s heteroplasmy (See **Suppl. Notes 1 and 2** for cautions). For example, (Otten et al. 2018) observed that levels of the 8993T>G mutation tended to increase in the offspring (positive HS). Some of us (Fleischmann et al. 2021), and then Zhang et al. (H. Zhang et al. 2021), also reported positive germline selection of 3243A>G, the most prevalent disease-causing mutation and a few other inherited pathogenic mutations.

The overall positive or negative selection does not tell the whole story, however. As illustrated in **Fig. 1**, the precise shape of the mutation’s “HS profile”, i.e. heteroplasmy shift as a function of heteroplasmy, critically affects the heteroplasmy changes with time and the mode of inheritance of a mutation and its impact on future generations and the population/species as a whole. The question of how germline selection in the germline depends of the heteroplasmy level has been addressed in several studies. Zhang et al. reported strongly *descending* HS profiles for several inherited pathogenic mutations (H. Zhang et al. 2021). In all cases, heteroplasmy shift was positive at *lower* heteroplasmy, and decreased, eventually turning negative, at higher heteroplasmy (e.g., the profile for 3243A>G mutations is shown in Fig 1A). Zhang et al. concluded that it was “likely that the positive selection at lower heteroplasmy levels counterbalanced the negative selection at higher levels”. We also have previously reported positive selection at low heteroplasmies and negative heteroplasmy shift correlation in m3243A>G, and proposed a similar, ‘naïve’ explanation at a conference (Khrapko 2019). This result, however, kept worrying us, primarily because the effect appeared unnaturally strong, and also the increase of the effect with decreasing heteroplasmy did not make biological sense. We, therefore, never published that observation is a journal and kept asking whether the apparent descending HS profile might have been a result of an error and sought to rectify the approach.

Analysis of the m3243A>G transmission requires compensating for life-long systematic changes of heteroplasmy in blood (Rajasimha, Chinnery, and Samuels 2008), (Grady et al. 2018). In hopes to improve the adjustment, we used the child’s heteroplasmy instead of the mother’s as a potentially more reliable reference because H_ch_ is expected to be developmentally ‘closer’ to the sought-after germline heteroplasmy and thus required a smaller and therefore more robust adjustment. This approach confirmed positive selection in the m3243A>G (Fleischmann et al. 2021), but, shockingly, the heteroplasmy shift profile was *inverted* and became ascending, i.e., the shift tended to be low/negative at low child’s heteroplasmy and to increase towards high child’s heteroplasmy. This apparent discrepancy was surprising because the child’s and mother’s heteroplasmies are correlated with each other, and from a biological standpoint, should not correlate with mtDNA transmission or selection in opposite ways. We therefore suspected that an artifact was skewing one or both approaches. Here we demonstrate that this is indeed an artifact i.e., the classical ‘regression to the mean’ (RTM) effect known for about 150 years (Galton 1886). We then show that RTM can be canceled and discuss the significance of the ‘true’ germline selection profile that becomes apparent after the removal of RTM (**Fig. 1**)

## Results

### The conventional negative correlation between the heteroplasmy shift and somatic heteroplasmy does not reflect the germline selection profile

As discussed in the Introduction, previous studies posed a controversy: the HS profile that, by the ‘naïve’ interpretation, is expected to represent the germline selection profile, appears to change from descending to ascending depending on whether mother’s (Khrapko 2019), (H. Zhang et al. 2021), or child’s (Fleischmann et al. 2021) heteroplasmy was used as the reference. This was alarming because it appeared to imply that HS cannot be used to measure of germline selection. Also, it called for an explanation of such an extraordinary discrepancy.

First, we confirmed the very existence of the discrepancy by repeating the mother-centered (HS(H_m_)) and the child-centered analyses (HS(H_ch_)) using exactly the same dataset. For this purpose we used one from (H. Zhang et al. 2021). The results of the resulting regression analyses are presented in **Fig. 2b and 2c**, respectively, and they fully confirm our concern: the two correlations are essentially symmetrical with opposite slopes. We also performed correlation analyses on other mutations for which extensive datasets are available, i.e., m8993T>G, m11778G>A, and m8344A>G using the data from (H. Zhang et al. 2021) for direct comparison with previous outcomes. The results are presented in supplementary **Fig. S2, a&d, b&e and c&f**, correspondingly. The results are qualitatively the same as for m3243A>G, which confirms that the observed opposite direction of slopes is a general phenomenon, not limited to the m3243A>G mutation.

**Figure 2.**
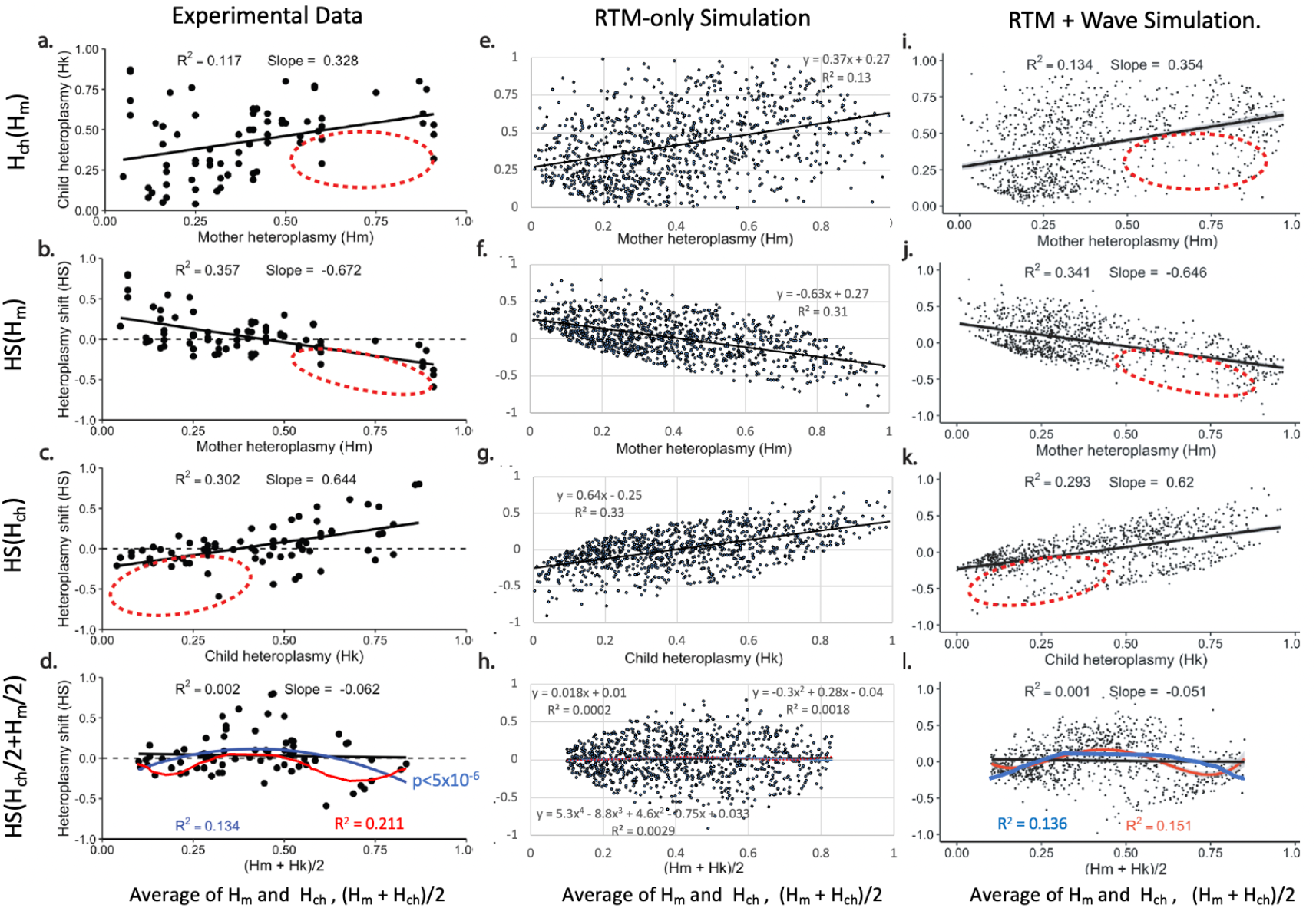
Regression analysis of experimenlal and simulated data demonstrates that germline selection follows a “wave-shaped profile”. **Data**. *Left column, (****a-d***): the experimental data - m3243A>G mother-child inheritance (public dataset, see methods for details). *Central column, (****e- h***): RTM-only simulation: Gaussian variance (SD=0.24) was added to the ‘exact transmission’ dataset {H_m_=H_ch_}, (0< H_m_,H_ch_<1). *Right column, (****i- l***): full simulation, i.e., exact transmission dataset with added Gaussian variance (0.135) AND the wave-shaped selection profile (see Methods for details). **Interpretation**: negative HS(H_m_) correlation (**b**) previously suggested that m3243A>G mutation was under germline selection that decreases from positive selection at low Hm, to negative selection at high Hm (see **Fig. 1A**). This hypothesis, however, is rejected, because it predicts that HS also decreases with Hk (Fig.S3 and discussion there), in contradiction with the observed positive correlation (**c**). Opposite correlations with H_m_ and H_ch_ are predicted if the data is affected by the “regression to the mean” bias (RTM), which is driven by data variance (**Fig. S1**). Indeed, our simulations show that experimentally observed correlations (**a-c**) may be fully accounted by the pure RTM (**e-g**). RTM Because RTM changes sign when H_k_ as a variable is changed to H_m_ (**f** vs. **g**), it follows that RTM can be nullified by the use of the symmetrical variable (H_m_+H_ch_)/2, as confirmed in (**h**). In support of the hypothesis that HS(H_m_) in experimental data is driven by RTM, this correlation was reduced to insignificant by the use of the (H_m_+H_ch_)/2 variable, as shown in (**d**). However, in the experimental dataset, cancellation of RTM revealed a highly significant (p<6×10^−5^) residual non-linear data skew, which is efficiently described by a quadratic (blue) or 4^th^-order (red) function (**d**, see also **Fig. S4**). This residual skew is, as expected, not present in the RTM-only simulation (**h**), which thus serves as an effective control. Accordingly, to advance the model, we complemented the RTM-only simulation with ‘wave-shaped selection’, i.e., the 4^th^ order function from (**d**); The RTM+Wave simulation (right column (**i** – **l))** correctly reproduces the residual 4^th^ order regression (**l)**. Moreover, it required the addition of only half of the variance needed for RTM-only simulation, because the added wave-shaped selection strengthened, synergistically with RTM, the HS(H_m_) and HS(H_ch_) correlations. Finally, RTM+Wave model reproduced the marked inhomogeneity of the experimentally observed datapoint ‘clouds’ (**a-c**). In particular, voids (red ellipses) in the simulated clouds appear reminiscent of the simlar irregularities in the experimental datapoint clouds. Statistical analysis of these similarities is complicated and is ongoing. We conclude that m3243A>G data distribution is mostly shaped by a combination of the wave-shaped germline selection and the RTM effect.

Next, we verified that the opposite HS(H_m_) and HS(H_ch_) slopes indeed rejected the ‘naïve’ interpretation of the descending HS(H_m_) profile as evidence of descending germline selection profile. To this end, we showed, both analytically and by a numerical simulation (Supplemental Fig. S3), that if germline selection profile were indeed descending, then heteroplasmy shift would have a strong *negative* component in the correlation *both* with mother’s *and* (even more so) with child’s heteroplasmy, which directly contradicts the observations in Fig. 2c and Fig. S2, and therefore decisively rejects the hypothesis that a descending HS profile means a descending germline selection profile.

### ‘Regression to the mean’ (RTM) drives both the negative HS(H_m_) and positive HS(H_ch_) correlation

In search of a phenomenon that could account for both negative HS(H_m_) correlation AND positive HS(H_ch_) correlation, we noted that negative correlations between HS and H_m_ in 3243A>G and other mutations (**Fig. 2d&g)** are highly reminiscent of the “regression towards mediocrity” phenomenon, first described in 1886 by Sir Francis Galton ((Galton 1886), *Fig. a* therein). This statistical bias, now called “regression towards mean” (RTM) refers to the tendency of parents who are ‘extreme’ with respect to a quantitative heritable trait (e.g., height) to have children who are closer to the population mean. The primary driver of the RTM effect is the non-inheritable variance of the trait. Extreme parents’ height is most likely due to a synergic combination of the extreme genetic and extreme non-genetic variance. The offspring most likely will lack the extreme non-genetic part, and thus will appear in most cases to converge to the population average (Barnett 2004). This tendency naturally results in a negative correlation between the level of the trait in the parents and the parent-to-child shift, just like we see with heteroplasmy shift. RTM is symmetrical with respect to the parent/child change. Indeed, if a child is extremely tall, then this is most likely due not only to genetics, but also to nongenetic factors that may not be shared with parents, so the parent(s) are likely to ‘regress to the mean’. Thus the RTM-driven correlation between HS and child heteroplasmy must be *positive*, i.e., opposite to that between HS and Hm, and exactly as observed in the real-world mtDNA inheritance data. Thus, RTM has the potential to explain the puzzling trends in the mtDNA inheritance.

First, we explored the behavior of the RTM effect on simulated data resembling the mtDNA inheritance dataset. We asked whether the RTM effect alone (i.e., RTM in the absence of germline selection) can account for the observed negative correlation between HS and H_m_. We started with the ‘exact transmission’ model (child inherits mother’s heteroplasmy level), in which H_ch_ is equal to H_m_ and both are uniformly distributed from 0 to 1. We then added gradually increased Gaussian variance to match the variance of the experimental data (see **‘Methods’, ‘RTM-only simulation’**). As discussed below in **Figure 3**, according to our biological interpretation, this variance is comprised of measurement error and variance generated in the somatic and germline bottlenecks. As standard deviation of the added variance in the simulation increased (0.1 to 0.3), the correlation between simulated H_ch_ and simulated H_m,_ expectedly, became weaker. The slope of H_ch_(H_m_) steadily decreased and the y-intercept increased (**Fig. S1**, top row, left to right). We thus used the comparison of the slope and the intercept to those of the experimental data as the criterion of the fitness of the model. The best fit variance (SD~0.24) was determined by comparing simulations at various SDs (see **Methods**). We then assessed the magnitude of the RTM effect in this ‘best-fit’ simulated dataset and compared it to the correlation of HS(H_m_) and HS(H_ch_) in the m3243A>G experimental dataset. Reassuringly, as shown in **Figure 2** (**b** vs. **f** and **c** vs. **g**), the two pairs of correlations (the simulated one and the experimentally observed one) are very similar. Thus, a realistic level of random variance can produce HS(H_m)_ and HS(H_ch_) correlations similar to those observed experimentally. Therefore, these observed HS(Hm) and HS(Hch) correlations may be entirely spurious, not related to selection. This discovery implies that nullification of the RTM bias may reveal the residual ‘true’ germline selection profile.

**Figure 3.**
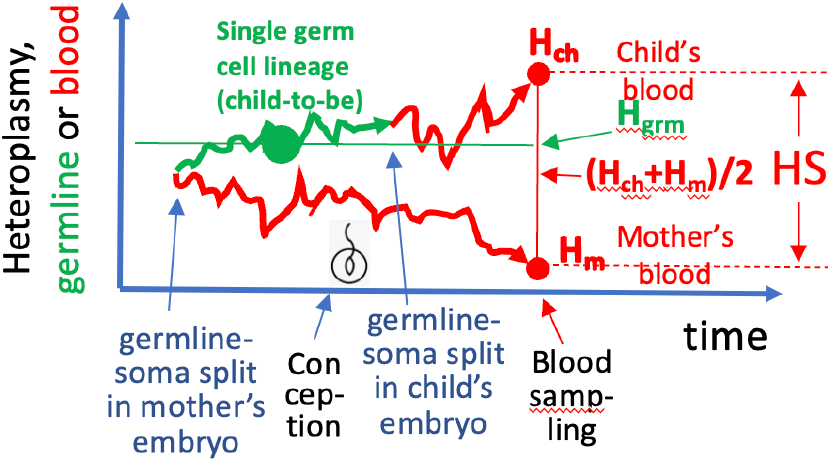
The dynamic of germline and blood heteroplasmy: (H_m_+H_c_)/2 is the optimal measure of germline heteroplasmy. The sketch shows qualitatively the constituents of the blood Heteroplasmy Shift (HS) of a hypothetical mother/child pair. The divergence of mother’s and child’s blood cells starts in mother’s embryo when mother’s somatic cells (including blood precursors) split from her germline. Red zig-zags illustrate the changes of the average heteroplasmy in the population of blood cells-to-be as they develop in the embryo and then in the blood. During these processes, heteroplasmy fluctuates stochastically through time and is eventually recorded as blood heteroplasmy ‘**H**_**m**_’ or ‘**H**_**ch**_’ at the time mother’s and child’s blood sampling, respectively. Note that the heteroplasmy trajectory of child’s blood first passes through mother’s germ line (green zig-zag). Unlike the multicellular blood trajectory, the germline trajectory follows a single germ cell lineage leading to the specific oocyte which becomes the child’s embryo after conception (fertilization symbol). If mother’s and child’s trajectories of blood heteroplasmy are random and unbiased then the best (on average) estimate of the heteroplasmy level in the germline (H_grm_) is the average of H_m_ and H_ch_, i.e., (H_m_+H_ch_)/2.

### Symmetrical (H_m_+H_ch_)/2 variable cancels RTM and reveals an underlying wave-shaped germline selection profile

As for nullification of RTM, we noted that because RTM-induced correlations have opposite signs depending on whether H_m_ or H_ch_ were used as the variable (**Fig. 2f** vs. **Fig. 2g**), they should cancel each other if the ‘symmetrical’ composite variable we used, such as (H_m_+H_ch_)/2. **Fig. 2h** confirms this prediction: there is no correlation between HS and (H_m_+H_ch_)/2 in the RTM-only model. Additional reason for the use of (H_m_+H_ch_)/2 is that it is more justified biologically than the conventionally used H_m_ measured in blood. Naturally, germline selection should depend on heteroplasmy in the germline, where selection takes place, not in the blood. As illustrated in **Fig. 3**, germline heteroplasmy is best approximated by (H_m_+H_ch_)/2) (also see Discussion). Thus, we used the (H_m_+H_ch_)/2) variable to cancel RTM and more appropriately represent the selection profile (**Fig. 2d**). Indeed, there is no significant linear trend in the experimental data plotted this way (**Fig. 2d**, black line). Intriguingly, we also noted an excess of positive HS measurements at intermediate values of (H_m_+H_ch_)/2. We therefore attempted to fit the data using quadratic regression, which indeed resulted in a highly significantly better fit (**Fig. 2d**, blue line). Reassuringly, there was no detectable quadratic component in the RTM-only simulated data (**Fig. 2h**, blue line). 4^th^ power regression (red in **Fig. 2d**) resulted increase of R^2^ compared to quadratic regression. Further increase of power to 5^th^ and 6^th^ added no additional improvement of the fitness (**Fig. S4**). We, therefore, chose 4^th^ power regression as our standard analysis of the HS((H_m_+H_ch_)/2) data plots. We postulate that this residual wave-shaped profile represents the true selection in the germline.

### Model that combines RTM and the wave-shaped selection precisely predicts experimental HS trends

The discovery of the residual wave-shaped selection in the experimental data implied that our RTM-only model that lacked such a residual (Fig. 2h) was incomplete. We therefore sought to incorporate the wave-shaped selection profile into the model. The ‘natural’ way to achieve this is to use an “input” selection curve to generate the simulated pairs of mother-child germline heteroplasmy dataset. To these selection-only data pairs, we added Gaussian variance in the same way as we did in the RTM- only simulation (see **Methods**). This approach is identical to the RTM-only simulation except here we used the selection-only data instead of the exact transmission data used in the RTM-only approach. To evaluate the model, we subjected the resulting simulated data to the regression analysis, identical to that used for experimental data and RTM-only simulation (Fig. 2). The criterion of success was (as in RTM-only) the similarity of the slopes and Y-intersects of regression lines of the simulated data to those of the m3243A>G experimental inheritance data (**Fig. 2**, left column). Additionally, we assessed if the model reproduces the the shape and amplitude of the observed residual wave-shaped curve (the “output” selection curve).

We started with a pilot simulation that used, as the input curve, intuitively, the experimentally observed m3243A>G residual wave-shaped 4^th^ power regression curve (**Fig. 2d)** and as the input variance, the variance of the best fit RTM-only model (Fig. 2i-l). This pilot simulation resulted in excessively steep regression slopes, both HS(H_m_) and HS(H_ch_) and excessively shallow slope of the Hk(Hm) correlation **(Suppl. Fig. S5 b, c, and a**, correspondingly). We concluded that (1) the addition of the selection into the model exacerbated the effect of RTM. Therefore the correlations in the experimental data must result from synergistic interaction between RTM and germline selection. At the same time, (2) the residual wave-shaped “output selection” curve was ‘dampened’, compared to the input curve (**Suppl. Fig. S5d**). This dampening effect is an expected result of added random noise. Thus, the real selection curve must be an augmented derivative of the experimentally observed residual curve shown in Fig. 2d.

Based on these two conclusions we optimized the model by (1) decreasing the added variance and (2) augmenting the added selection curve. We performed several iterations of optimization which greatly improved the simulation. Reassuringly, optimization simultaneously improved the fit along all the criteria: the HS(H_m_) and HS(H_ch_) slopes and Y-intersections, the R^2^ of correlations, and the shape of the ‘output’ residual selection curve, and eventually precisely satisfied all optimization criteria.

The results of the optimized simulation are shown in **Fig. 2i-l** (right column). Additionally, simulation appears to reproduce the otherwise mysterious ‘voids’ in the datapoint clouds (shown by ellipsoids in **Fig. 2**). Furthermore, the RTM-plus-selection simulation accounted for otherwise puzzling disparity between HS(H_m_) and HS(H_ch_) slopes, which under no conditions could be reproduced in RTM-only simulations (see **Supplemental Methods** for more detail). Whereas the statistical significance of ‘the voids’ and ‘the disparity’ is difficult to assess, their mere presence collectively boosts our confidence in the simulations. We conclude that the wave-shaped selection profile model fully accounts for the key features of the m3243A>G dataset which supports the hypothesis that the germline selection of these mutations indeed follows such a profile. As a note of caution, our analysis has not excluded the possibility that other models could equally reproduce the dataset’s properties.

### Wave-shaped selection is a general phenomenon: analysis of other disease-causing mtDNA mutations

To explore the generality of the wave-shaped selection profile, we performed regression analysis of other human mtDNA mutations for which rich enough datasets are available: m8993T>G, m11778G>A, and m8344A>G using data from Zhang et al. to ensure compatibility of the results. The analyses are shown in the Supplement, **Fig. S2**. Similar to m3243A>G and as expected, all mutations show strong negative HS(H_m_) correlations (**Fig. S2, a-c**). Also, similarly to m3243A>G and to the RTM-only simulation, the correlation of HS(H_ch_) is positive (**Fig. S2, d-f**), which suggests the presence of the RTM effect. And, finally, m8993T>G and m11778G>A also show a wave-shaped selection profile when plotted against (H_m_+H_ch_)/2 (**Fig. S2, g, h**). Additionally, both of these mutations show overall more positive germline selection than m3243A>G (average HS across all heteroplasmy levels is more positive). Interestingly the last mutation, m8344A>G does not show an arch, but a weak monotonous *increase* of HS with (H_m_+H_ch_)/2 (**Fig. S2i**). This is the opposite of the inference for this mutation using H_m_ as the independent variable.

This difference between m8344A>G and other three mutations may imply fundamental dissimilarities between the mechanisms of germline selection. From methodological point, this observation reassures us that the arched shape observed in other mutations is not an artifact inherent to our analysis, but a true biological phenomenon. This is effectively a negative control in the experimental data, which adds to the negative control in the simulated data without selection (i.e. **Fig. 2h**), though more data are needed for definitive conclusions.

## Discussion

### Possible mechanisms underlying the ‘wave-shaped’ selection profile

It is tempting to speculate regarding the possibe mechanism(s) responsible for the ‘wave-shaped’ selection profile which was uncovered by removing the RTM effect. It appears likely that the complexity of the profile is a result of a superposition of multiple processes, some of which are responsible for purifying selection, and some for positive selection. Indeed, there is substantial literature both on positive (Holt, Speijer, and Kirkwood 2014; Kowald and Kirkwood 2018) and negative (Gottlieb et al. 2021) selection of mitochondria and mitochondrial mutations. Importantly, purifying selection can proceed on a single mitochondrion level, for example, by a mitochondrion- autonomous negative feedback loop such as the PINK1-Larp loop (Y. Zhang et al. 2019). This implies that purifying selection may be operational at very low mutant fractions, which potentially accounts for the prevalence of negative selection in the left extreme of the selection profile. There are very little data available with low H_m_ values, however, and definitive conclusions about germline selection at low heteroplasmy would be premature.

Purifying negative selection on a single mitochondrion level mentioned above (Y. Zhang et al. 2019) may be expected to counteract positive selection that we observe at higher heteroplasmy levels. This effect, however, appears to be not strong enough as detrimental mtDNA mutations are known to be under positive selection in tissues as diverse as muscle (Nicholas et al. 2009), choroid plexus (Campbell et al. 2012) and colon (Taylor et al. 2003). One therefore must assume that mitochondrion-autonomous purifying selection may either cease at higher mutant fractions and/or be overcome by positive selection. This assumption fits well with the wave-shaped selection profile, though specific mechanisms explaining positive selection are still a matter of debate (Holt, Speijer, and Kirkwood 2014), (Kowald and Kirkwood 2018). Note that at intermediate heteroplasmy levels, positive selection does not necessarily need to proceed at a single mitochondrion level. As heteroplasmy levels exceed physiological threshold (Konstantin Khrapko and Turnbull 2014), the physiology of the entire cell may change, which, for example, may make the cell more prone to proliferation (Smith et al. 2020). Finally, at even higher mutant fractions, (as has been repeatedly proposed by many researchers in the field) detrimental mutations may become progressively more toxic to the cell and cause slowdown of proliferation or active removal of highly mutant germ cells cells (e.g. by attrition (Fan et al. 2008)), or by embryo demise.

### Biological justification of the use of the (H_m_+H_ch_)/2 variable

In this study we used the symmetrical (H_m_+H_ch_)/2 variable to cancel the RTM effect. There is another compelling reason to use (H_m_+H_ch_)/2 as an independent variable instead of the H_m_, which is traditionally used to describe germline selection of mtDNA. Note that there is no consensus regarding the precise mechanism and location of mtDNA bottleneck, which is currently variably described by several competing models (Cao et al. 2007; Cree et al. 2008; K. Khrapko 2008; Lee et al. 2012; Wai, Teoli, and Shoubridge 2008). Similarly, there is little consensus about the location, mechanism, and even direction of germline selection. Nevertheless, most models (Fleishmann et al. 2021; Floros et al. 2018; H. Zhang et al. 2021) postulate that selection takes place sometime during and/or after the embryonic mtDNA bottleneck (Cree et al. 2008). Granted, selection process must be controlled not by the mother’s heteroplasmy in blood (H_m_), but by heteroplasmy in germ cells around the bottleneck. As schematically illustrated in **Fig. 3**, blood heteroplasmy H_m_ is separated from the ‘around-bottleneck’ heteroplasmy (large green dot, ‘H_grm_’) by changes in heteroplasmy during blood development and the lifelong cell dynamics in the blood (Rajasimha, Chinnery, and Samuels 2008), (Franco et al. 2022; Grady et al. 2018), which collectively represent a ‘somatic bottleneck’ (Barrett et al. 2020). Given that these random changes take place in blood samples from both mothers and children, (H_m_+H_ch_)/2 must be a better proxy of heteroplasmy level at the point where selection is expected to take place than the conventionally used mother’s heteroplasmy in blood.

### Biological significance of the wave-shaped selection profile

Negative or zero selection in the lower heteroplasmy segment (left extreme of the wave, **Fig. 1**) is expected to prevent nascent mutations (which are initially necessarily occur at a very low fraction) from expansion. Most of them are destined to disappear in the nearest cell generations, largely due to intracellular random genetic drift (Coller, Bodyak, and Khrapko 2002). However, drift is expected to push a small proportion of nascent mutations to intermediate fractions. These mutations potentially pose the danger of being spread into subsequent generations and further into the population. With the wave selection profile, these mutations will be further enriched to even higher levels by the positive selection that prevails at intermediate heteroplasmy levels. The resulting individuals (or oocytes, or embryos) with a detrimental mutation at high heteroplasmy levels will likely be unfit and therefore removed from population by Darwinian selection (or oocyte attrition, or embryo demise), before mutation is spread further. Our preliminary simulations imply that this strategy may indeed be efficient in protecting the genetic pool of the population under realistic assumptions (in preparation). In contrast, with the continuously descending selection profile (grey line in **Fig. 1A**), positive selection at low heteroplasmy will increase the proportion of nascent mutations that reach higher fractions, which means increased mutational pressure. Furthermore, mutations that reach intermediate fractions are not pushed to highly detrimental levels. Instead, they are expected to converge to intermediate level both from below and from above. Thus, they are expected to remain at mildly detrimental levels which are most dangerous as far as transmission to offspring and contaminating of the population’s genetic material is concerned.

### Clinical significance

Our findings have important clinical implications. On a general note, a significant role of RTM implies that random processes strongly impact the predictions of the transmission of pathogenic mitochondrial mutations. We demonstrate the advantage of using a proxy of the actual germline heteroplasmy, the average between rather than child and mother heteroplasmy in blood. In the clinical setting, the task usually entails the prediction of child heteroplasmy based on the mother’s, so the child’s heteroplasmy is not known and it is not possible to use the (H_m_+H_ch_)/2 variable. However, a better estimate of the germline heteroplasmy level can be achieved by averaging the heteroplasmy levels of the developmentally distant tissues, such as blood, buccal cells, urine epithelial cells – all these are relatively easy to collect from the patient. Our analysis suggests that such a combined assessment of the germline heteroplasmy levels may potentially significantly improve the predictions for prospective children of individuals carrying pathogenic mtDNA mutations.

### Future directions

We note that mitigation of RTM by using the (H_m_+H_ch_)/2 variable is just the first, though likely a major, step in elucidating of the selection profiles based on mother/child heteroplasmy shift data. One of the results of this study was the identification of RTM as a major bias in analysis of mtDNA inheritance and to find a way to factor it out. We recognize, however, that additional, finer, adjustments may be needed to establish the selection profiles with higher accuracy. We anticipate that two aspects need to be addressed. The HS-based selection analysis considers absolute changes of heteroplasmy, whereas relative changes better reflect the “selection intensity”. Consideration of relative changes is particularly important at low heteroplasmy levels, where absolute changes are small and uninformative. However, relative heteroplasmy changes become unstable at low heteroplasmy, because of small numbers is in denominator. Small random variance thus may result in huge chandes as as We hope thatb eventually the availability of larger numbers of high quality data will allow precise measurents of selection in the vicinity of zero heteroplasmy.

## Methods

### Regression analysis

To perform the analyses of the effect of regression to the mean, for each of the experimental and simulated mother-child inheritance datasets (i.e. any collection of (H_m_,H_ch_) pairs of (mother, child) heteroplasmy measurements) we first calculated, for every mother-child pair, heteroplasmy shift HS= H_ch_ - H_m_, and plotted heteroplasmy shifts as a function of mother’s heteroplasmy HS(H_m_), child heteroplasmy HS(H_ch_), and also as a function of the ‘symmetrized’, average mother/child heteroplasmy, HS((H_m_+H_ch_)/2). Linear and polynomial regression analysis of the data was performed using Excel’s built-in analysis tools.

***Note***: *all heteroplasmies used in this figure (Hm, Hk) are measured or simulated as heteroplasmies in blood. The graphs in the last row, however, represent heteroplasmy shift in the germline because (H*_*m*_*+H*_*ch*_*)/2 represents heteroplasmy in the germline*.

### RTM-only simulation (Fig. 2 middle column, and Fig. S1)

In the RTM-only simulation is based on the exact transmission model, i.e. we assumed that the child inherits its mother’s heteroplasmy without change (H_m_ = H_ch_). This model ensures the absence of any systematic selection and no variance. Specifically, we started with 10,000 pairs of equal values (H_m_ = H_ch_) sampled uniformly between 0 and 1 (i.e. sampled with a 0.0001 step). To emulate the RTM effect, we added variance to this model by adding to each mother and child heteroplasmy value a number randomly drawn from a Gaussian distribution. We performed a series of such simulations using various standard deviation values to define the Gaussian distribution (the results of simulations with SD=0 (the original undisturbed model) and datasets with added Gaussian noise with SD=0.1, SD=0.2, and SD=0.3 are shown in **Fig. S1**. Data pairs in which one or both of the simulated mother/child heteroplasmy values was outside the 0.1-0.85 range were deleted. Note that for fairness of comparison, such mother/child pairs were also deleted from the experimental 3243 Zhang dataset.

*Additionally, to avoid potential biases related to the difference between the uniform distribution of the simulated data and the uneven distribution of experimental data points we emulated the experimental distribution of data points in our simulations and picked the simulated data points from the experimental distribution*.

As expected, the slope and R^2^ of the linear regression between H_m_ and H_ch_ systematically decreased as standard deviation of the added Gaussian variance was increased. At the same time, also as expected, the strength of the RTM effect (represented by the slopes of linear regressions of the distributions HS(H_m_) and HS(H_ch_)) increased in absolute terms, but in opposite directions (**Fig. S1** second and third row, correspondingly). We chose the slope and R2 of the linear regression between H_ch_ and H_m_ as a criterion to select the simulation that best fitted the experimental m3243A>G dataset. The addition of variance using a Gaussian distribution with a standard deviation of 0.275 ensured best fit (in the simulation a slope of 0.35 and R^2^ of 0.12, vs a slope of 0.33 and R^2^ of 0.12 in the experimental data – see **Fig. 2a** and **Fig. 2b**). We then subjected the simulated dataset with an SD of 0.275 to the ‘RTM analysis’ described above and compared the result to that of the Zhang dataset of 3243 mutation inheritance. The resulting plots show close similarity between the experimental and simulated datasets with respect to all linear correlations (**Fig. 2**, compare plots on the left column to those in the center column). Figure 2 (center column) is a representative subsample of the simulated dataset, which was obtained by down-sampling the simulation to 85 datapoints for easy visual comparison to the experimental data. The regression parameters shown in center column pertain to the full simulation (10,000 datapoints).

### RTM+Wave simulation (Fig. 2, right column)

To assess the combined effect of wave-shaped germline selection and RTM, we first generated a 10,000 – strong set of data pairs representing the systematic selection in the germline. This dataset consisted of {H_m_+H_c_)/2, HS((H_m_+H_c_)/2)} pairs of values which followed an appropriate wave-shaped curve (e.g., the 4^th^ order curve in **Fig. 2d**, or adjusted curves in the optimization simulations). According to our scenario, this systematic heteroplasmy change in the germline was then succeeded with random variance as heteroplasmic mtDNA was distributed into the blood cells (**Fig. 3**). Consequently, we first converted the germline heteroplasmy level H_m_+H_c_)/2 into the corresponding heteroplasmy levels in mother and child, {H_m_, H_ch_} and then added the appropriate random variance to both H_m_ and H_ch_. Such conversion is possible by solving a system of linear equations for each {H_m_+H_c_)/2, HS((H_m_+H_c_)/2)} pair. Then the resulting ‘noisy’ dataset was subject to the regression analysis shown in Fig 2 and described in the ‘regression analysis’ section above.

In the first pilot simulation, we used as a first approximation the input parameters the experimentally observed residual arched profile (Fig 2d, red curve) and variance of the RTM-only simulation (**Fig. 2e-h**). The first pilot simulation, resulted in excessively strong HS(H_m_) correlation in the simulated data (the regression lines, both HS(H_m_) and HS(H_ch_) were steeper than in the experimental dataset. Also, the output residual regression curve was ‘dampened’, i.e. whereas the overall shape of the curve was unchanged, the minima and the maxima were reduced about 1.5-fold. The mechanism of this effect is similar to the deterioration of a sharp signal peak in a noisy environment. Thus the residual regression curve of the experimental m3243A>G in **Fig. 2d** is a blunted derivation of the real germline selection profile. We therefore concluded that to reconstruct the original selection profile and to achieve a high-fit simulation, the standard deviation of the added variance must be reduced and the input wave-shaped curve must be augmented (i.e., the absolute values of the maximum and the minima must be proportionally increased compared to the experimentally observed residual curve). With that in mind, we performed several iterations of the simulations and determined that decreasing the standard deviation of the added variance to 0.21 and augmenting the wave-shaped curve

1.5 times ensures an optimized fit of the summary features of the m3243A>G experimental data. The results of the optimized simulation are presented in **Fig. 2**, right column.

### Monte Carlo estimation of significance

To determine whether the non-linear selection profile was a statistically significant finding, we determined the probability that the quadratic regression of the magnitude observed in the experimental m3243A>G data appears by pure chance in the absence of selection. We performed 100,000 random samplings of 85 datapoints each (to match the 85 ndataspoints in the experimental set) from the RTM-only simulated dataset and then counted the number of instances where the R^2^ of the second order best-fit quadratic curve exceeded 0.134, which is the quadratic R^2^ of the experimental m3243A>G dataset. There were six such cases, putting the p-value below 6×10^−5^.

## Abbreviations/conventions

H_m_: *heteroplasmy in mother’s blood*,
H_ch_: *heteroplasmy in child’s blood;*
HS: *H*_*ch*_ **–** *H*_*m*_ *heteroplasmy shift*.
HS(H_m_): heteroplasmy shift as a function of mother’s heteroplasmy. The syntax in HS (H_ch_), HS((*H*_*m*_*+H*_*ch*_*)/2*), and *H*_*ch*_ (H_m_) carries the equivalent meaning.

## Data availability

In this study we used the mtDNA mutation inheritance datasets published by Zhang et al., for the purpose of consistency. In this dataset, the m3243A>G data was corrected for the decay of heteroplasmy levels in blood with age.

## Notes

### Competing Interest Statement

The authors have declared no competing interest.

### Summary of Updates

The revision incorporates further analysis of the source of the RTM artifact, including novel modeling work.

## References

Barnett, A. G. 2004. “Regression to the Mean: What It Is and How to Deal with It.” International Journal of Epidemiology 34 (1): 215–20. 10.1093/ije/dyh299.

Barrett, Alison, Barbara Arbeithuber, Arslan Zaidi, Peter Wilton, Ian M. Paul, Rasmus Nielsen, and Kateryna D. Makova. 2020. “Pronounced Somatic Bottleneck in Mitochondrial DNA of Human Hair.” Philosophical Transactions of the Royal Society B: Biological Sciences 375 (1790): 20190175. 10.1098/rstb.2019.0175.

Campbell, G. R., Y. Kraytsberg, K. J. Krishnan, N. Ohno, I. Ziabreva, A. Reeve, B. D. Trapp, et al. 2012. “Clonally Expanded Mitochondrial DNA Deletions within the Choroid Plexus in Multiple Sclerosis.” Acta Neuropathologica 124 (2): 209–20. 10.1007/s00401-012-1001-9.

Cao, L., H. Shitara, T. Horii, Y. Nagao, H. Imai, K. Abe, T. Hara, J. Hayashi, and H. Yonekawa. 2007. “The Mitochondrial Bottleneck Occurs without Reduction of mtDNA Content in Female Mouse Germ Cells.” Nature Genet 39 (3): 386–90.

Coller, H. A., N. D. Bodyak, and K. Khrapko. 2002. “Frequent Intracellular Clonal Expansions of Somatic mtDNA Mutations: Significance and Mechanisms.” Ann N Y Acad Sci 959 (April): 434–47.

Cree, L. M., D. C. Samuels, S. C. de Sousa Lopes, H. K. Rajasimha, P. Wonnapinij, J. R. Mann, H. H. Dahl, and P. F. Chinnery. 2008. “A Reduction of Mitochondrial DNA Molecules during Embryogenesis Explains the Rapid Segregation of Genotypes.” Nature Genet 40 (2): 249–54.

Fan, W., K. G. Waymire, N. Narula, P. Li, C. Rocher, P. E. Coskun, M. A. Vannan, J. Narula, G. R. MacGregor, and D. C. Wallace. 2008. “A Mouse Model of Mitochondrial Disease Reveals Germline Selection against Severe mtDNA Mutations.” Science 319 (5865): 958–62.

Fleischmann, Zoe, Sarah J. Pickett, Melissa Franco, Dylan Aidlen, Mark Khrapko, David Stein, Natasha Markuzon, et al. 2021. “Bi-Phasic Dynamics of the Mitochondrial DNA Mutation m.3243A&gt;G in Blood: An Unbiased, Mutation Level-Dependent Model Implies Positive Selection in the Germline.” bioRxiv, January, 2021.02.26.433045. 10.1101/2021.02.26.433045.

Fleishmann, Zoe, Sarah J. Pickett, Melissa Franco, Dylan Aidlen, Mark Khrapko, David Stein, Natasha Markuzon, et al. 2021. “Bi-Phasic Dynamics of the Mitochondrial DNA Mutation m.3243A&gt;G in Blood: An Unbiased, Mutation Level-Dependent Model Implies Positive Selection in the Germline.” bioRxiv. 10.1101/2021.02.26.433045.

Floros, Vasileios I., Angela Pyle, Sabine Dietmann, Wei Wei, Walfred C. W. Tang, Naoko Irie, Brendan Payne, et al. 2018. “Segregation of Mitochondrial DNA Heteroplasmy through a Developmental Genetic Bottleneck in Human Embryos.” Nature Cell Biology 20 (2): 144–51.

Franco, Melissa, Sarah J Pickett, Zoe Fleischmann, Mark Khrapko, Auden Cote-L’Heureux, Dylan Aidlen, David Stein, et al. 2022. “Dynamics of the Most Common Pathogenic mtDNA Variant m.3243A>G Demonstrate Frequency-Dependency in Blood and Positive Selection in the Germline.” Human Molecular Genetics 31 (23): 4075–86. 10.1093/hmg/ddac149.

Galton, Francis. 1886. “Regression Towards Mediocrity in Hereditary Stature.” The Journal of the Anthropological Institute of Great Britain and Ireland 15: 246. 10.2307/2841583.

Gottlieb, Roberta A., Honit Piplani, Jon Sin, Savannah Sawaged, Syed M. Hamid, David J. Taylor, and Juliana de Freitas Germano. 2021. “At the Heart of Mitochondrial Quality Control: Many Roads to the Top.” Cellular and Molecular Life Sciences 78 (8): 3791–3801. 10.1007/s00018-021-03772-3.

Grady, John P., Sarah J. Pickett, Yi Shiau Ng, Charlotte L. Alston, Emma L. Blakely, Steven A. Hardy, Catherine L. Feeney, et al. 2018. “mtDNA Heteroplasmy Level and Copy Number Indicate Disease Burden in m.3243A&gt;G Mitochondrial Disease.” EMBO Molecular Medicine 10 (6): e8262. 10.15252/emmm.201708262.

Holt, Ian J., Dave Speijer, and Thomas B. L. Kirkwood. 2014. “The Road to Rack and Ruin: Selecting Deleterious Mitochondrial DNA Variants.” Philos Trans R Soc Lond B Biol Sci 369 (1646).

Khrapko. 2019. “MtDNA Bottleneck against Dertrimental mtDNA Mutations: What Does It Do, and What It Does’t.” In. Moscow, Russia.

Khrapko, K. 2008. “Two Ways to Make an mtDNA Bottleneck.” Nature Genet 40 (2): 134–35.

Khrapko, Konstantin, and Doug Turnbull. 2014. “Mitochondrial DNA Mutations in Aging.” Progress in Molecular Biology and Translational Science 127: 29–62. 10.1016/b978-0-12-394625-6.00002-7.

Kowald, Axel, and Thomas B. L. Kirkwood. 2018. “Resolving the Enigma of the Clonal Expansion of mtDNA Deletions.” Genes 9 (3): 126. 10.3390/genes9030126.

Lee, H. S., H. Ma, R. C. Juanes, M. Tachibana, M. Sparman, J. Woodward, C. Ramsey, et al. 2012. “Rapid Mitochondrial DNA Segregation in Primate Preimplantation Embryos Precedes Somatic and Germline Bottleneck.” Cell Reports 1 (5): 506–15. 10.1016/j.celrep.2012.03.011.

Nicholas, A., Y. Kraytsberg, X. Guo, and K. Khrapko. 2009. “On the Timing and the Extent of Clonal Expansion of mtDNA Deletions: Evidence from Single-Molecule PCR.” Exp Neurol 218 (2): 316–19.

Otten, Auke B. C., Suzanne C. E. H. Sallevelt, Phillippa J. Carling, Joseph C. F. M. Dreesen, Marion Drüsedau, Sabine Spierts, Aimee D. C. Paulussen, et al. 2018. “Mutation-Specific Effects in Germline Transmission of Pathogenic mtDNA Variants.” Hum Reprod 33 (7): 1331–41.

Rajasimha, Harsha Karur, Patrick F. Chinnery, and David C. Samuels. 2008. “Selection against Pathogenic mtDNA Mutations in a Stem Cell Population Leads to the Loss of the 3243A→G Mutation in Blood.” The American Journal of Human Genetics 82 (2): 333–43. 10.1016/j.ajhg.2007.10.007.

Smith, Anna L. M., Julia C. Whitehall, Carla Bradshaw, David Gay, Fiona Robertson, Alasdair P. Blain, Gavin Hudson, et al. 2020. “Age-Associated Mitochondrial DNA Mutations Cause Metabolic Remodeling That Contributes to Accelerated Intestinal Tumorigenesis.” Nature Cancer 1 (10): 976–89.

Stewart, James Bruce, Christoph Freyer, Joanna L. Elson, and Nils-Goran Larsson. 2008. “Purifying Selection of mtDNA and Its Implications for Understanding Evolution and Mitochondrial Disease.” Nat Rev Genet 9 (9): 657–62. 10.1038/nrg2396.

Taylor, R. W., M. J. Barron, G. M. Borthwick, A. Gospel, P. F. Chinnery, D. C. Samuels, G. A. Taylor, et al. 2003. “Mitochondrial DNA Mutations in Human Colonic Crypt Stem Cells.” J Clin Invest 112 (9): 1351–60.

Wai, T., D. Teoli, and E. A. Shoubridge. 2008. “The Mitochondrial DNA Genetic Bottleneck Results from Replication of a Subpopulation of Genomes.” Nature Genet 40 (12): 1484–88. 10.1038/ng.258.

Wonnapinij, Passorn, Patrick F. Chinnery, and David C. Samuels. 2010. “Previous Estimates of Mitochondrial DNA Mutation Level Variance Did Not Account for Sampling Error: Comparing the mtDNA Genetic Bottleneck in Mice and Humans.” The American Journal of Human Genetics 86 (4): 540–50.

Zhang, Haixin, Marco Esposito, Mikael G. Pezet, Juvid Aryaman, Wei Wei, Florian Klimm, Claudia Calabrese, et al. 2021. “Mitochondrial DNA Heteroplasmy Is Modulated during Oocyte Development Propagating Mutation Transmission.” Science Advances 7 (50): eabi5657. 10.1126/sciadv.abi5657.

Zhang, Yi, Zong-Heng Wang, Yi Liu, Yong Chen, Nuo Sun, Marjan Gucek, Fan Zhang, and Hong Xu. 2019. “PINK1 Inhibits Local Protein Synthesis to Limit Transmission of Deleterious Mitochondrial DNA Mutations.” Molecular Cell 73 (6): 1127-1137.e5. 10.1016/j.molcel.2019.01.013.

